# Explainable Fuzzy Clustering Framework Reveals Divergent Default Mode Network Connectivity Dynamics in Schizophrenia

**DOI:** 10.1101/2023.02.13.528329

**Authors:** Charles A. Ellis, Robyn L. Miller, Vince D. Calhoun

## Abstract

Dynamic functional network connectivity (dFNC) analysis of resting state functional magnetic resonance imaging data has yielded insights into many neurological and neuropsychiatric disorders. A common dFNC analysis approach uses hard clustering methods like k-means clustering to assign samples to states that summarize network dynamics. However, hard clustering methods obscure network dynamics by assuming (1) that all samples within a cluster are equally like their assigned centroids and (2) that samples closer to one another in the data space than to their centroids are well-represented by their centroids. In addition, it can be hard to compare subjects, as in some cases an individual may not manifest a state strongly enough to enter a hard cluster. Approaches that allow a dimensional approach to connectivity patterns (e.g., fuzzy clustering) can mitigate these issues. In this study, we present an explainable fuzzy clustering framework by combining fuzzy c-means clustering with several explainability metrics. We apply our framework for schizophrenia (SZ) default mode network analysis, identifying 5 states and characterizing those states with a new explainability approach. While also showing that features typically used in hard clustering can be extracted in our framework, we present a variety of unique features to quantify state dynamics and identify effects of SZ upon network dynamics. We further uncover relationships between symptom severity and interactions of the precuneus with the anterior and posterior cingulate cortex. Given the ease of implementing our framework and its enhanced insight into network dynamics, it has great potential for use in future dFNC studies.

## INTRODUCTION

Resting state functional magnetic resonance imaging (rs-fMRI) dynamic functional network connectivity (dFNC) data has historically been used to give insight into a variety of neurological [1] and neuropsychiatric disorders [2]–[6] and cognitive functions [7][8]. A common dFNC analysis approach involves applying a hard clustering approach (e.g., k-means clustering) to assign dFNC samples to a set of dynamical states that are supposedly representative of the overall time series [2], [9]–[15]. Features can then be extracted based on the identified time-resolved states that can give insight into various aspects of the state dynamics. This method has been widely applied but makes a critical assumption that could obscure useful disease-related dynamics. Namely, it assumes that all samples assigned to a state equally belong to a state. This is problematic given that samples very near to one another in the data space may be assigned to two distant cluster centroids. A few studies have presented fuzzy clustering approaches that indicate the degree to which samples belong to different states, but this area is still highly understudied [11], [15]. Although explainability methods have been developed specifically to help characterize identified states, the use of hard clustering methods can make explainability difficult. In this study, we present a novel explainable fuzzy clustering framework for fMRI dFNC that identifies fuzzy states and assigns samples a probability of belonging to each state. We further present novel dynamical features that use the output probabilities and demonstrate their utility by applying them to identify differences in the dynamics of default mode network (DMN) activity between individuals with schizophrenia (SZs) and healthy controls (HCs).

Several modalities have been used for insight into the effects of neurological and neuropsychiatric disorders upon brain dynamics. These include electroencephalography (EEG), magnetoencephalography (MEG), and fMRI. All three modalities – EEG [16]–[18], MEG [19], and fMRI [13], [20]–[24] - have been used extensively for SZ analysis. EEG and MEG capture much higher resolution temporal information. However, localizing the region of the brain associated with MEG and EEG signals can be challenging. In contrast, fMRI has much higher spatial resolution at a lower temporal resolution relative to EEG and fMRI. A select group of studies have also combined multimodal EEG, MEG, or fMRI data for insight into disorders [9]. However, in general, fMRI is more often used in SZ analysis than the other modalities. Within fMRI analysis, both task [25], [26] and resting state [10], [12], [13], [27]–[32] data are frequently used. However, resting state data offers several advantages. Specifically, the majority of brain activity is spontaneous (i.e., better captured by resting state), so resting state analysis provides an avenue to understand how the brain operates under most circumstances [32]. Additionally, task performance in healthy controls relative to individuals with schizophrenia often varies greatly, introducing a potential confounder into any neuroimaging analyses [32].

Many studies have analyzed resting state fMRI data within the context of SZ and other neuropsychiatric disorders. For example, studies have used independent components (ICs) [24], spectral features [9][33], and functional network connectivity. Functional network connectivity offers a unique benefit in that it provides insights into the interaction of different brain regions and networks. Early in the use of rs-fMRI functional network connectivity, it was more common to analyze the correlation between brain regions across a whole recording. This approach is referred to as static functional network connectivity (sFNC) [34]. Although sFNC had widespread use, a number of studies found that dFNC (i.e., functional connectivity captured in windows over time) offered insights into brain interactions that would otherwise be obscured by sFNC analysis [32], [34]. Both sFNC and dFNC have been used to gain insight into a variety of neurological and neuropsychiatric disorders and cognitive functions, including Alzheimer’s disease [1], [35], [36], major depressive disorder [3]–[6], schizophrenia [2][37], cognition [7], and spatial orientation [8]. However, dFNC offers greater opportunities to learn about the brain than sFNC. As discussed in [32], [34], early studies extracted functional network connectivity from time-series extracted using one seed per brain region, multiple seeds from a given region of interest (ROI), multiple seeds from subregions of multiple ROIs, and seeds from whole-brain ROIs. However, in recent years, the fully automated, group independent component analysis (ICA)-based Neuromark pipeline has been developed as a approach for extracting time-series used to calculate functional network connectivity [38]. It yields components that are reproducible across datasets and studies and contains components from a variety of brain networks and subregions. Furthermore, it has been used in many studies [1], [3], [24], [35]–[42].

Multiple approaches have been used to analyze dFNC extracted using the NeuroMark pipeline or other approaches. Many studies have used classification approaches [21], [22], [43] or a combination of clustering and classification [39]. However, a described in [32], a many studies have used clustering approaches [9]–[15]. These clustering approaches involve assigning dFNC samples to states that summarize the dFNC time-series. Features can then be extracted to quantify various aspects of the state time-series. By far the most common clustering approach used for identifying states of dFNC activity is k-means clustering [44]. K-means clustering involves randomly initializing cluster centroids, calculating the average of the samples nearest each centroid, updating the cluster centroids to be equal to the average of the nearest samples, and iteratively repeating the process up to the point that the cluster centroid stops moving at each iteration above a pre-defined threshold. The main advantage of k-means clustering for use with dFNC is that it is easy to implement using existing libraries [45]. However, while k-means clustering is widely used, it does have some disadvantages. First, it can yield low quality clusters. Previous studies have proposed approaches to try to address this problem [14]. Additionally, it does not assign each sample a probability of belonging to each cluster centroid or resting-state, which involves an implicit assumption in subsequent analysis that the identified states are able to adequately summarize the dFNC time-series [46]. This is a big assumption given that samples very near one another in the data space can be assigned to completely different states. In addition, in the hard clustering approach it can be hard to compare individuals as it is likely that some individuals may never enter a given state strongly enough to enter the cluster. As such, a simple fuzzy clustering-based approach could go a long way towards helping future studies in the field of dFNC clustering to uncover new aspects of disorder-related brain dynamics by better summarizing brain activity.

As the field of rs-fMRI FNC clustering has developed, a few studies have introduced explainability approaches with the goal of helping characterize the differences between identified states [7], [23], [47]. In contrast to earlier efforts involving statistical hypothesis testing [38], these methods quantify the importance of FNC features in a manner that acknowledges the multi-dimensional nature of the underlying clustering. However, these methods have an important shortcoming that arises from their use with hard clustering methods. Specifically, they perturb FNC features and examine the sensitivity of the underlying clusters to perturbation. They quantify the effect of this perturbation upon the clustering by calculating the percentage of samples that completely switch clusters. Given that only a small minority of samples switch clusters following perturbation, it is safer to say that the methods quantify the effects of perturbation upon that subset of samples rather than upon the overall clustering. Similar to how a fuzzy clustering approach could help future studies better summarize brain activity and uncover new aspects of disorders, fuzzy clustering could also contribute to the development of explainability approaches that are better able to quantify the effects of perturbation.

In this study, we present fuzzy c-means clustering [48] as an approach for the identification of fMRI dFNC fuzzy states. Fuzzy c-means has been used widely across application areas like emotion recognition from speech data [49], customer segmentation [50], and brain image segmentation [51]. Furthermore, fuzzy c-means is simple to use given its inclusion in several publicly available Python packages [52] and MATLAB [53]. As such, it has the potential to be widely used within the domain of FNC clustering. Our approach outputs a probability that each sample belongs to each fuzzy state. After performing clustering, we present a pair of new explainability approaches to characterize the identified states, comparing them to an existing approach [47]. We then present a variety of novel dynamical and stability features that highlight different aspects of the fuzzy state time-series while also showing that our approach is compatible with dynamical features that have been used in previous hard clustering studies. We further show how the features can be used to differentiate between healthy controls and individuals with schizophrenia in the default mode network (DMN), a network that has previously been associated with SZ [10], [12], [13], [27]–[32], and identify relationships between the features and SZ and SZ symptom severity.

## METHODS

In this section, we describe and discuss our approach for the study. Figure 1 presents an overview of our approach. (1) We used a pre-existing rs-fMRI dataset composed of individuals with schizophrenia (SZs) and healthy controls (HCs). (2) We preprocessed the data and extracted dFNC. (3) We applied fuzzy c-means clustering to identify 5 fuzzy states. (4) We sought to identify the important dFNC features for each state by applying and comparing the results for two global clustering explainability methods. (5) We next extracted a number of pre-existing and novel dynamical features to summarize different aspects of the identified states. (6) We applied a novel local cluster explainability approach for insight into the stability of study participants to the perturbation of specific dFNC features. (7) To identify SZ-related differences in the extracted dynamical and stability features, we conducted statistical analyses and trained a series of logistic regression classifiers with elastic net regularization (LR-ENR). (8) Lastly, we performed statistical analyses to determine whether the dynamical and stability features were related to symptom severity.

**Figure 1.**
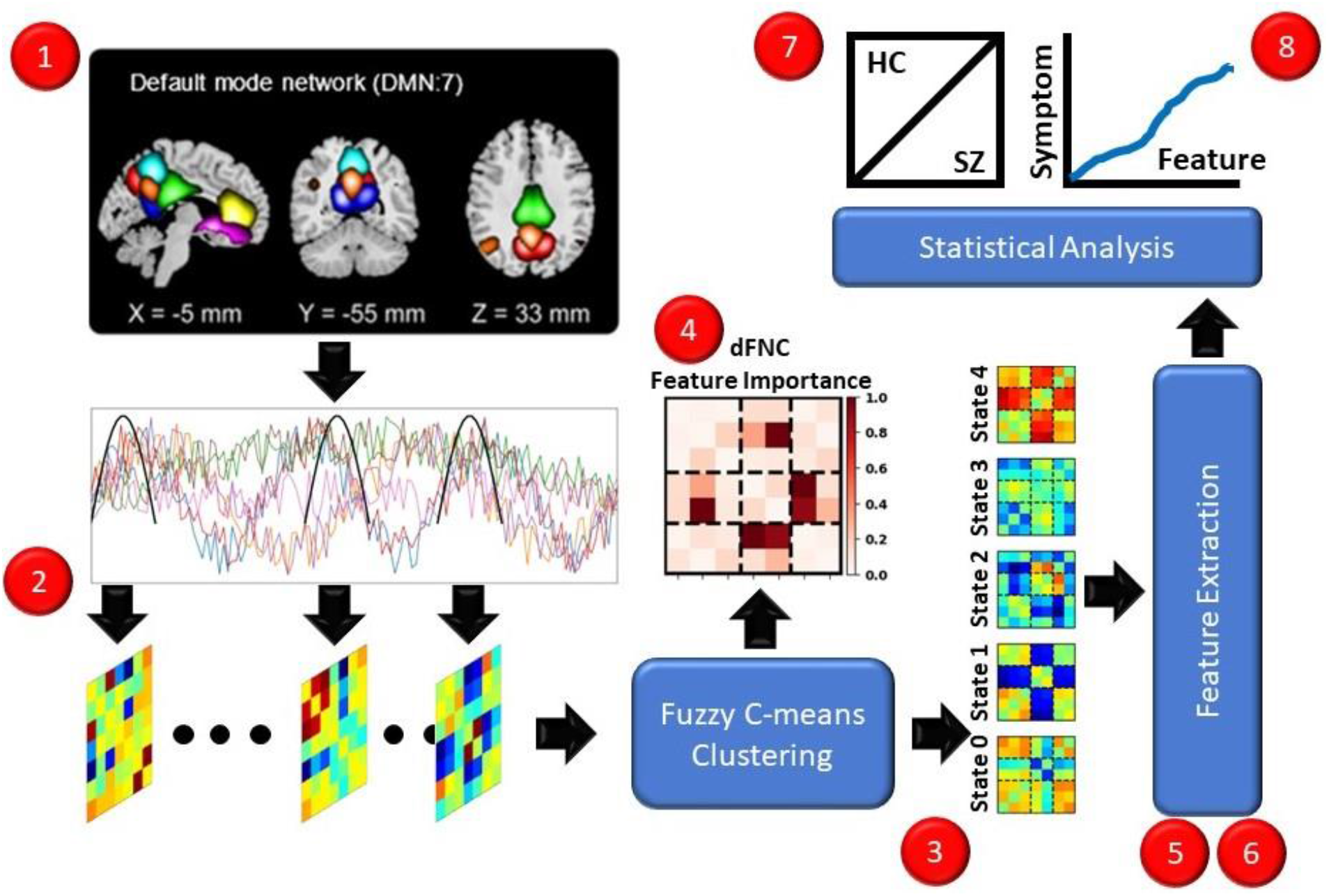
Overview of Methods. Red dots indicate each step of the methods. (1) We recorded rs-fMRI data from HCs and SZs. (2) We extracted dFNC data. (3) We performed fuzzy c-means clustering, identifying 5 states. (4) We applied several explainability approaches for insight into the dFNC features characterizing the states. (5,6) We extracted dynamical and stability features. (7) We performed t-tests and trained interpretable machine learning models for insight into the features that differed between HCs and SZs. (8) We performed linear regression analyses controlling for age and gender to identify relationships between symptom severity and dynamical and stability features.

### Description of Dataset

In this study, we used the Functional Imaging Biomedical Informatics Research Network (FBIRN) dataset, consisting of rs-fMRI recordings from 151 SZs and 160 HCs [54]. The dataset has been used in many studies, both related to fMRI dFNC clustering and classification [21], [24], [43], [47]. In addition to neuroimaging data, positive and negative symptom severity scores from the Positive and Negative Syndrome Scale (PANSS) [55]. Positive symptoms of SZ include hallucinations, delusions, and bizarre behavior. Negative symptoms include alogia, apathy, affective flattening, and asociality [28]. The dataset was collected at 7 sites: the University of California at Los Angeles, the University of California at San Francisco, the University of California at Irvine, the University of Iowa, the University of Minnesota, the University of New Mexico, and Duke University/the University of North Carolina at Chapel Hill. Data collection procedures were approved by the institutional review boards of each center, and all study participants gave written informed consent. One site used a 3T GE MR750 scanner, and 6 sites used 3T Siemens TIM Trio Scanners. Data was collected using a T2*-weighted AC-PC aligned EPI sequence (TE = 30ms, TR = 2s, slice gap = 1mm, flip angle = 77°, voxel size = 3.4 × 3.4 × 3.4 mm^3^, acquisition time = 5 min and 38s, and number of frames = 162).

### Description of Data Preprocessing

Prior to preprocessing, the first 5 mock scans were removed. Statistical parametric mapping (SPM12, https://www.fil.ion.ucl.ac.uk/spm/) was used for preprocessing, and head motion was corrected using rigid body motion correction. The data was spatially normalized to an echo-planar imaging template in the standard Montreal Neurological Institute (MNI) space. Following resampling to 3×3×3 mm^3^, a Gaussian kernel with a 6mm full width at half maximum was used to smooth the data. The fully automated Neuromark Pipeline of the Group ICA of fMRI Toolbox (GIFT, http://trendscenter.org/software/gift) involved the use of spatially constrained ICA to extract corresponding components while adaptive to individual datasets. In our case, we used the neuromark_fMRI_1.0 template to extract 53 independent components (ICs) with peak activations in the gray matter of various brain networks. Seven of the ICs were associated with the DMN, and we used those components in this study. The 7 ICs included 3 precuneus (PCN), 2 anterior cingulate cortex (ACC), and 2 posterior cingulate cortex (PCC). After IC extraction, dFNC was estimated using Pearson’s correlation with a sliding tapered window. The window consisted of a rectangle with a 40-second step size convolved with a Gaussian (σ=3). While a number of step sizes have been utilized in previous studies, a 40-second step size is highly common [13], [23], [37], [56]. The dFNC extraction resulted in a dataset composed of 21 dFNC features and 124 time steps per participant. During the remainder of this paper, correlation between two brain regions will be abbreviated using the pattern of IC1/IC2 (e.g., PCC1/ACC2), where IC1/IC2 is equivalent to IC2/IC1.

### Description of Clustering Approach

After extracting dFNC, we concatenated all dFNC time steps across samples and applied fuzzy c-means clustering [48]. Fuzzy c-means clustering is comparable to k-means clustering, but unlike k-means clustering, fuzzy c-means assigns each sample a probability of belonging to a given cluster. We used the Python package scikit-fuzzy in our implementation [52]. We initialized the clustering with 100 random seeds and selected the seed from the initialization with the highest fuzzy partitioning coefficient. It is relatively common to set the number of clusters to a number identified in previous studies [57]. As such, to make it easier to compare our findings to previous studies [13] of DMN dFNC data in SZ, we used 5 clusters. A number of other studies have also found 5 clusters to be ideal for clustering SZ dFNC data [57]. We used a maximum number of iterations of 1000 and an error of 0.0001. Additionally, we optimized the fuzziness parameter, m, selecting from potential values of 1.01, 1.5, and 2.

### Description of Explainability Approach

We used three global explainability approaches for insight into the relative importance of each dFNC feature to the identified fuzzy states. We used Global Permutation Percent Change (G2PC) feature importance [47]– an existing approach that has been used in several neuroimaging studies [7], [23], and we used two variations of a novel approach called, “Permutation-based Distribution Divergence (P2D)”. There are global (GP2D) and local (LP2D) variations of P2D based upon how the divergence estimates for individual samples are consolidated. After applying each approach, we compared them.

### G2PC

G2PC is an adaptation of permutation feature importance to the domain of unsupervised clustering explainability. Permutation feature importance was originally developed to explain random forest models [58] and was later extended in a model-agnostic manner to explain a variety of supervised models [59]. G2PC involves (1) permuting a particular feature across all samples in a dataset for a given number of repeats, (2) reassigning the perturbed samples to the previously identified clusters, and (3) calculating the percentage of samples in each repeat that switch clusters. Feature importance is considered to linearly increase with the percentage of samples that switch clusters following the perturbation of a feature (i.e., increased sensitivity of the cluster to a feature corresponds to increased importance of that feature). In our analysis, we permute each feature 1,000 times.

### GP2D

A key shortcoming of G2PC is that in a data space with a high number of dimensions perturbing a single feature typically only affects a minority of samples. The use of fuzzy c-means clustering, rather than a hard clustering method like k-means clustering, presents a new opportunity. Specifically, the sensitivity of the clustering to perturbation can be measured as the change in the distribution of probabilities for each sample belonging to each cluster, rather than the percentage of samples that completely switch clusters. As such, our novel approach involves the same first two steps as G2PC. However, we quantify the effect of perturbation by calculating the Kullback-Leibler divergence (KLD) [60] between the original probabilities of a given sample belonging to each cluster and the probabilities for the perturbed sample belonging to each cluster. We repeated the permutation for 1,000 repeats and then calculated two summary metrics from the KLD. (1) We calculated the median KLD across all samples for each repeat (i.e., median GP2D), and (2) we summed the total KLD across all samples for each repeat (i.e., total GP2D).

### G2PC and GP2D Comparison

We hypothesized that the total GP2D would be more likely to be strongly affected by the select group of samples that would be sensitive to G2PC, and that the median GP2D metric would differ from both G2PC and total GP2D by accounting for more of the distribution of the effects of perturbation across all samples. To test these hypotheses, we ranked each feature based on its median importance across all 1,000 repeats for each method, producing 3 rankings. We then applied Kendall’s rank correlation [61] in SciPy [62] in a pair-wise approach, comparing G2PC to total GP2D, G2PC to median GP2D, and total GP2D to median GP2D. After obtaining p-values for each test, we applied FDR correction [63] to reduce the likelihood of false positives.

### LP2D

Building upon the GP2D approach, we also calculated the mean of the KLD for each sample across repeats. We used the resulting values for two sets of analyses. (1) We wanted to determine whether our novel P2D approach was actually able to capture the effects of perturbation upon more samples than G2PC, so we calculated the percentage of samples that had non-zero KLD values. (2) We wanted to use LP2D as an approach for estimating the stability of individuals to each dFNC feature. To that end, we also used the standard deviation of the mean of LP2D values across samples for each participant as dynamical features (i.e., for each sample take the mean KLD across repeats and then take the standard deviation of the resulting values across samples belonging to each subject). These stability features and the specific analyses performed upon them will be further discussed in subsequent sections.

### Description of Dynamical Feature Extraction

After clustering all of the samples, we extracted a number of features to quantify various aspects of the dynamics of the transition to and from the identified fuzzy states. We used two types of features that have been frequently applied in previous studies involving hard clustering and developed a number of new features uniquely suited for use with fuzzy clustering that give new insight into state dynamics. We sought to use these features to understand the effects of SZ upon the DMN dynamics, and statistical analyses using the features will be described in subsequent sections.

### Traditional Features

We extracted two types of traditional features: the occupancy rate (OCR) and number of state transitions (NST). These features have been used in many previous studies. Extracting these features required thresholding the probabilities of each sample for each fuzzy cluster such that each sample was assigned to the cluster for which it had the highest probability of belonging. The OCR is the percentage of time points that each participant spends in each state, so with 5 dFNC states there are 5 OCRs for each participant. The NST is the number of times that each participant changes states.

### KLD and Entropy-based Features

The first set of novel dynamical features that we introduced used Kullback-Leibler divergence and Shannon entropy. For the KLD-based features, we calculated the KLD of the 5 fuzzy state probabilities between each of the 124 consecutive time steps for each participant. This yielded an array of KLD values between each time point with which we calculated a number of additional values. For each participant, we calculated the mean KLD across time points, the median KLD across time points, the maximum KLD between any two consecutive time points, the range of KLD values between time points, and the KLD between probabilities for the first and last time points in a recording. The KLD-based features quantified different aspects of how the distribution of probabilities across fuzzy states shifted over time. However, we also wanted to gain insight into how the probabilities of each individual fuzzy state shifted over time. To this end, we calculated the Shannon entropy across all time points for each fuzzy states separately (i.e., yielding one entropy value per state).

### Descriptive Statistic-based Features

We next used a number of traditional descriptive statistics as novel dynamical features. Specifically, for each state, we calculated the mean, variance, and range of probabilities across time points for each participant. This yielded 15 features (i.e., 1 mean, 1 variance, and 1 range feature per state). The mean value gave an approximation of how strongly a particular participant resides in a particular fuzzy state. Similar to the Shannon entropy feature, the variance value gives insight into how much the similarity of the dFNC features for each participant varies over time. Lastly, whereas the mean feature would capture how strongly a participant resided within a given state and the variance feature would capture how the similarity of a participant to each fuzzy state varied over time, the range feature sought to give insight into the more extreme probabilities for each time-series that might otherwise be obscured. The utility of this feature makes sense given that previous studies have identified the effects of SZ upon the brain to be highly localized [24].

### Correlation-based Features

Lastly, we wanted to identify relationships between specific states. As such, we calculated the correlation between the probabilities for each state over time. This yielded 10 features. The use of correlation could give insight into whether probabilities shifting from a particular state over time are redistributed to a specific alternative state or redistributed to a number of states.

### Cumulative Difference Features

Similar to how previous studies have sought to understand the total distance traveled within a particular state [11], we wanted to understand in an absolute manner how much the probabilities of a participant being in a state change over time. As such, we calculated the difference between probabilities for each state across each pair of consecutive time points and summed the absolute differences for each state. This yielded 5 features (i.e., 1 feature per state).

### Uniformity Feature

Lastly, we sought to quantify the degree to which state probabilities were uniformly distributed across states. To this end, we calculated the absolute difference between the 5 state probabilities at each step and the uniform probability of 0.2. We then summed the absolute differences at each step and averaged across time steps for each participant.

### Description of Class-Related Statistical Analysis

We wanted to determine whether the fuzzy states that we identified were related to SZ dynamics. As such, we performed a series of two-tailed t-tests comparing the dynamical and LP2D stability features belonging to SZs to those belonging to HCs. After performing the t-tests, we performed separate FDR corrections on the OCR features, correlation-based features, cumulative change features, and LP2D-based features to reduce the likelihood of false positive test results.

### Description of LR-ENR Classification-based Explainability Analysis

Our statistical analysis indicated whether there were significant differences in the extracted dynamical features between HCs and SZs and gave insight into the effects of SZ upon DMN dynamics. However, we also wanted to determine how helpful the extracted features could be for discriminating between HCs and SZs and gain insight into the relative importance of each feature for the classification. To this end, we classified SZs and HCs using LR-ENR. LR-ENR is a frequently used [13] interpretable machine learning model. Its coefficients can be visualized to obtain insight into the relative importance of each feature included in the classification. We feature-wise z-scored each feature and then trained separate LR-ENR classifiers for the traditional, non-uniformity + KLD + entropy, variance, mean, range, correlation, cumulative difference, and LP2D features. We used 10-fold nested cross-validation with 64%, 16%, and 20% of the data being assigned to training, validation, and test sets, respectively. Within the inner cross-validation folds, we optimized the ratio of L1 to L2 normalization, selecting from values of 0.25, 0.5, 0.75, 0.85, 0.9, 0.95, and 0.99. We further optimized the inverse of the regularization strength, selecting from 100 values geometrically spaced between 10^−4^ and 10^4^. We used the saga solver from Scikit-Learn [45], and allowed a maximum of 200,000 iterations. After obtaining the optimal parameter set for each inner fold, we retrained each model on both training and validation data using the optimal parameters before testing. We then calculated the area-under-curve (AUC) of the receiver-operating-curve to quantify the performance of each model. We calculated the mean and standard deviation of the AUC, sensitivity (SENS, i.e., true positive rate), and specificity (SPEC, i.e., true negative rate) for the 10 folds corresponding to each set of dynamical features.

### Description of Symptom Severity Analysis

Lastly, while our earlier analyses gave insight into differences in DMN dynamics between SZs and HCs, we also wanted to determine whether the fuzzy states we uncovered could be used for insight into SZ symptom severity. As such, we performed ordinary least squares regression using age, gender, and the negative PANSS score or age, gender, and the positive PANSS score as independent variables and the dynamical features as independent variables. We lastly applied FDR correction for the OCR features, entropy features, correlation-based features, cumulative change features, and LP2D-based features separately.

## RESULTS

In this section, we describe the 5 fuzzy states that we identified with our clustering approach. We then show the important dFNC features identified using our global explainability analyses. After characterizing the states and the dFNC features that differentiate them, we examine whether there are differences in the dynamical features and LP2D-based stability features between HCs and SZs and whether the features are related to symptom severity.

### Identifying 5 Fuzzy States of dFNC Activity

As shown in Figure 2, we identified 5 distinct fuzzy states of dFNC activity. State 0 was the only state to have highly negative intra-ACC dFNC. It had moderately highly levels of positive PCC/PCN and intra-PCC dFNC and a mixture of low magnitude positive and negative dFNC for ACC/PCN and ACC/PCC, where ACC1 and ACC2 had slightly positive and negative dFNC, respectively. State 1 was characterized as having highly negative ACC/PCN and ACC/PCC dFNC and almost no PCN/PCC and intra-region dFNC. Relative to the other states, state 2 had more varied patterns of positive and negative dFNC. PCC1/PCN dFNC was highly negative, while PCC2/PCN dFNC was a mixture of low magnitude positive and negative activity. ACC/PCN2 and ACC/PCN3 were highly or moderately negative, but ACC/PCN1 had almost no connectivity. Intra-region dFNC was mostly low magnitude with the exception of intra-PCC dFNC, which was highly negative. State 3 was the only state to have negative intra-PCN dFNC. Additionally, although most other regions had slightly positive dFNC, PCC2/PCC and intra-PCC activity had low to moderately negative dFNC. Lastly, state 4 contrasted heavily with state 1 with very high positive ACC/PCN and ACC/PCC activity. State 4 also had low to moderately positive dFNC between all other pairs of regions.

**Figure 2.**
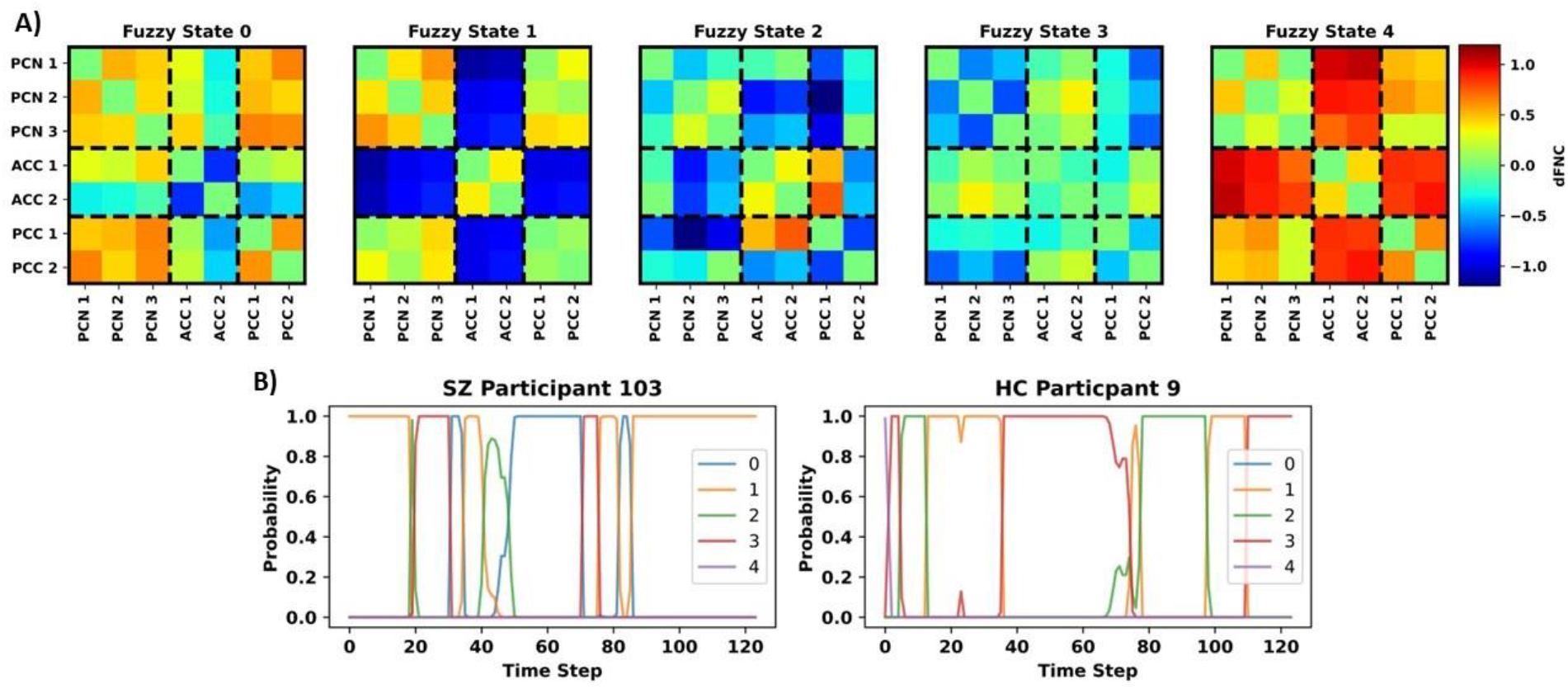
Fuzzy States and Example State Trajectories. Panel A shows the centroids for each of the 5 fuzzy states that we identified. Each subpanel shares the same color bar to the right of the panel for Fuzzy State 4. The ICs associated with each dFNC feature are arranged on the x- and y-axes and are grouped based upon brain region (i.e., PCN, ACC, PCN). Panel B shows example state trajectories for an SZ participant (left) and HC participant (right). The y-axis indicates the probability of belonging to each state, and the different colors of lines correspond to the state numbers shown in the legends. Note the variation in probability of belonging to each cluster.

### Identifying Key dFNC Features Differentiating Each State and Comparing Explainability Approaches

Although we visually compared the centroids of the clusters that we identified, the visual comparison did not address the relative importance of each dFNC feature to the identified states in a quantitative manner and did not consider the underlying distribution of the samples in relation to the identified fuzzy states. Figure 3 shows the results for our global explainability analyses. G2PC, total GP2D, and median GP2D all identified ACC/PCC1 and ACC/PCN2 to be among the most important dFNC features with the other features being of varying importance. G2PC affected around 10% of samples. In contrast, mean LP2D captured the effects of perturbation on 100% of samples. Additionally, there was some high-level agreement in the relative importance (i.e., importance rank) of each dFNC feature across global approaches (e.g., the most important features for one method tended to be among the most important for the other methods). However, there was not a statistically significant Kendall’s rank correlation between the global feature importance across methods. G2PC and total GP2D had a correlation coefficient of -0.048. G2PC and median GP2D and a correlation coefficient of 0.229, and total GP2D and median GP2D had a correlation coefficient of 0. Additionally, while G2PC and total GP2D tended to distribute importance widely, median GP2D provided a much sparser importance estimate.

**Figure 3.**
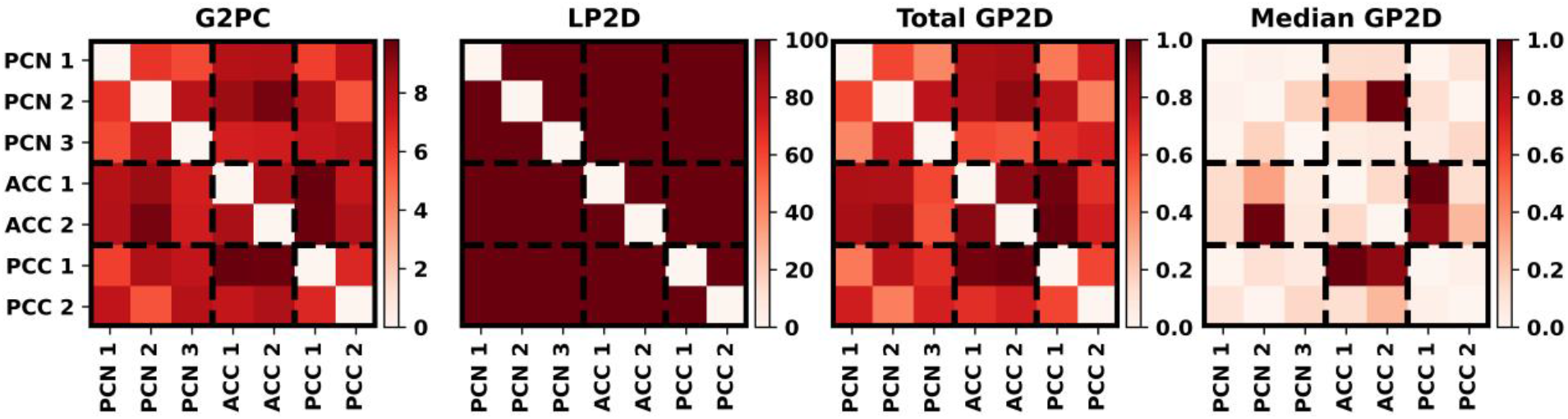
Explainability Results. From left to right, the panels show the G2PC, LP2D, total GP2D, and median GP2D results. The color bar corresponding to each panel is positioned to its right, and the ICs associated with each dFNC feature are arranged on the x- and y-axes and are grouped based upon brain region (i.e., PCN, ACC, PCN). Total GP2D and median GP2D are scaled by their maximal value for easier interpretation.

### Identifying Disorder-Related Differences in Dynamical and Stability Features

Figure 4 shows the results for our t-tests examining differences in dynamical and LP2D stability features between HCs and SZs, and Supplementary Figures 1 through 4 show boxplots of the dynamical features for each class. Many features had statistically significant differences between groups. SZs had significantly higher values in four of the six KLD-based features (i.e., max, range, SD, KLD). Although SZs had a slightly but not significantly higher mean KLD relative to HCs, there was almost no difference in their median KLD relative to HCs. SZs had significantly higher state 0 entropy and lower state 2 entropy. Interestingly, SZs also had higher and lower state 0 and state 2 OCRs, average probabilities, variance in probabilities, ranges of probabilities, and cumulative differences, respectively. Additionally, state 0/1 and 0/3 correlation had a more negative magnitude in SZs than HCs, while state 0/2, state 1/2, state 2/3, and state 2/4 SZs had a less negative magnitude than HCs. Lastly, SZs had higher levels of sensitivity to PCC2/PCN1, PCC2/PCN3, and intra-ACC perturbation than HCS and lower levels of sensitivity to PCC1/PCN2 perturbation than HCs.

**Figure 4.**
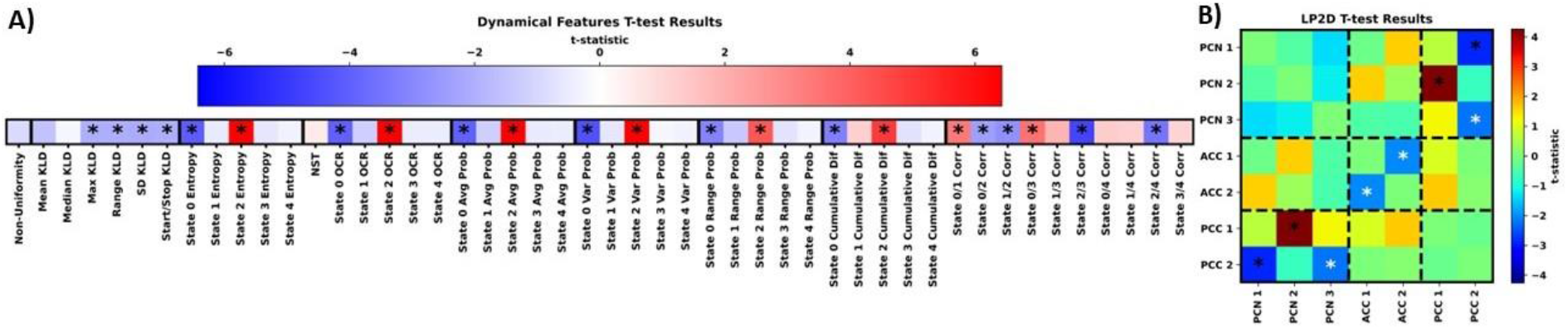
Group-Level Statistical Comparison. Panels A and B show the t-test results for the dynamical and LP2D stability features, respectively. In panel A, the groups of features we extracted are arranged from left to right, and dark black lines separate each group of features. In panel B, results are arranged in the form of a standard connectivity matrix for easier interpretation. Dashed lines separate domain pairs. Both panels are heatmaps showing the t-statistics for each feature. The color bar for panel A is above the heatmap, and the color bar for panel B is to the right of the heatmap. Black and white asterisks indicate features with significant differences with and without FDR correction, respectively It should be noted that the t-tests were performed as HCs minus SZs. As such, a negative t-statistic indicates that SZs had higher values for a particular feature than HCs.

Table 2 shows the LR-ENR performance results, and Figure 5 shows the LR-ENR explainability results. As might be expected, given the t-test results, the NST + OCR, Non-Uniformity + KLD + Entropy, Average, Variance, Range, and Correlation feature models had the highest AUC values. The LP2D and cumulative difference feature models had slightly lower performance. The Range model had the highest overall AUC. For SENS, the NST + OCR and Average feature models performed highest, with Variance feature model performing slightly lower. For SPEC, the Range, and Non-Uniformity + KLD + Entropy models performed highest. The model coefficients also provided results highly consistent with the t-test results, exhibiting similar directionality and relative magnitude of differences between HCs and SZs.

**Table 1.** Description of Study Participants

|  | Number | Age | Gender (M/F) | PANSS (positive) | PANSS (negative) |
| --- | --- | --- | --- | --- | --- |
| **SZ** | 151 | 38.06 ± 11.30 | 115/36 | 15.32 ± 04.92 | 14.32 ± 05.42 |
| **HC** | 160 | 37.04 ± 10.68 | 115/45 | not applicable | not applicable |

**Table 2.** LR-ENR Performance Results

| Features | AUC | SENS | SPEC |
| --- | --- | --- | --- |
| **NST + OCR** | 68.58 ± 06.61 | 70.00 ± 06.46 | 59.06 ± 06.32 |
| **Non-Uniformity + KLD + Entropy** | 67.77 ± 06.47 | 56.77 ± 06.80 | 65.63 ± 06.09 |
| **Average** | 68.59 ± 06.77 | 68.06 ± 05.29 | 60.31 ± 08.73 |
| **Variance** | 67.72 ± 07.24 | 63.23 ± 07.80 | 63.44 ± 08.03 |
| **Range** | 69.18 ± 06.50 | 43.23 ± 06.49 | 74.69 ± 07.32 |
| **Correlation** | 66.05 ± 06.18 | 55.81 ± 08.17 | 65.00 ± 08.59 |
| **Cumulative Difference** | 62.24 ± 05.33 | 54.19 ± 10.03 | 61.25 ± 05.26 |
| **LP2D** | 60.56 ± 09.41 | 51.61 ± 08.16 | 60.00 ± 09.56 |

**Figure 5.**
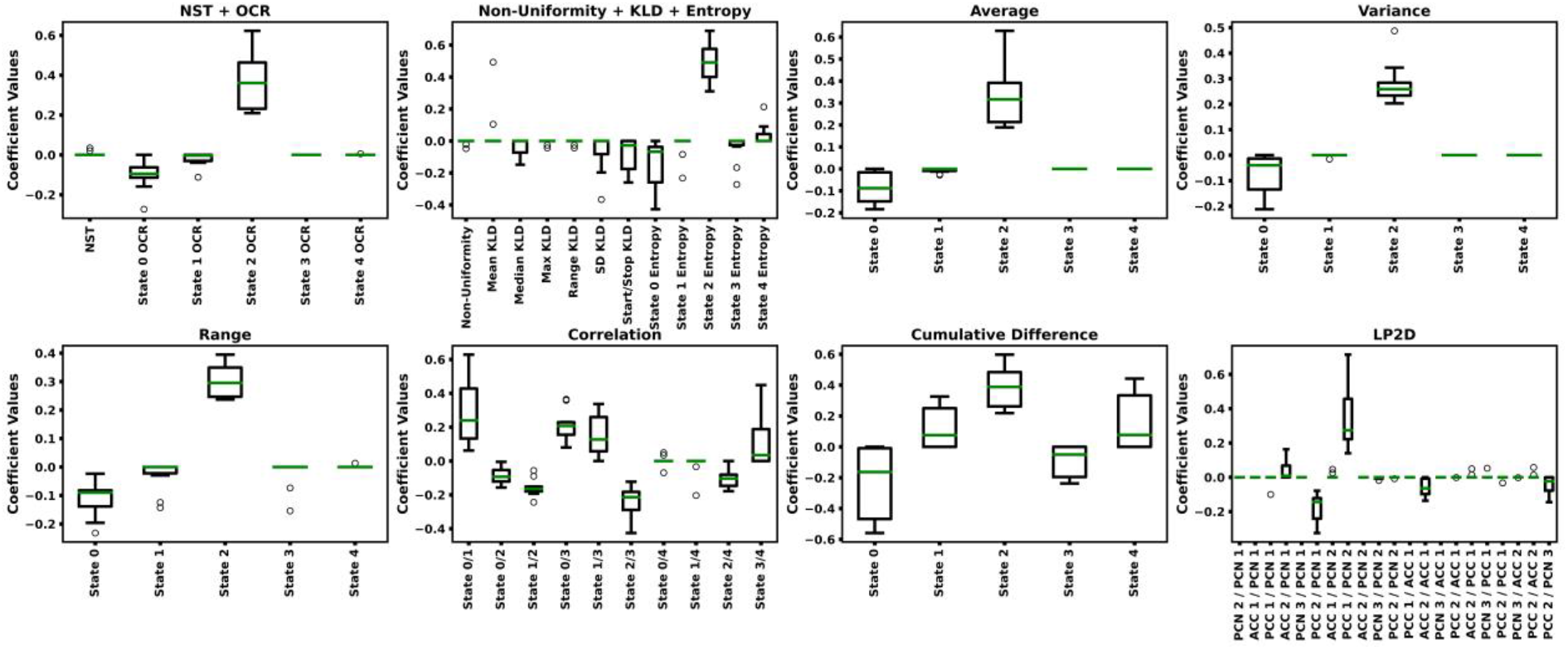
LR-ENR Explainability Results. Each panels shows the LR-ENR coefficient values for the groups of features indicated in their respective titles. Specific feature names are indicated on the x-axis of each panel. Coefficient values are shown on the y-axis. Higher magnitude coefficients indicate greater importance to their respective model. Because a label of 0 was used for SZs and a label of 1 was used for HCs, a negative coefficient value indicates that an increase in that feature corresponded to an increase in likelihood of belonging to the SZ class.

### Identifying Relationship between Symptom Severity and Dynamical and Stability Features

Interestingly, no dynamical features had significant relationships with or without FDR correction with symptom severity, so we did not include a figure detailing the results for that analysis. However, as shown in Figure 6, without FDR correction multiple LP2D features had significant relationships with positive and negative symptom severity. Increased sensitivity to PCC1/PCN2 and PCC1/PCN3 perturbation corresponded to increases in both positive and negative symptom severity. Additionally, increased sensitivity to ACC1/PCN2 perturbation corresponded to increased positive symptom severity, and increased sensitivity to ACC1/PCN1 and intra-PCC perturbation corresponded to increased negative symptom severity.

**Figure 6.**
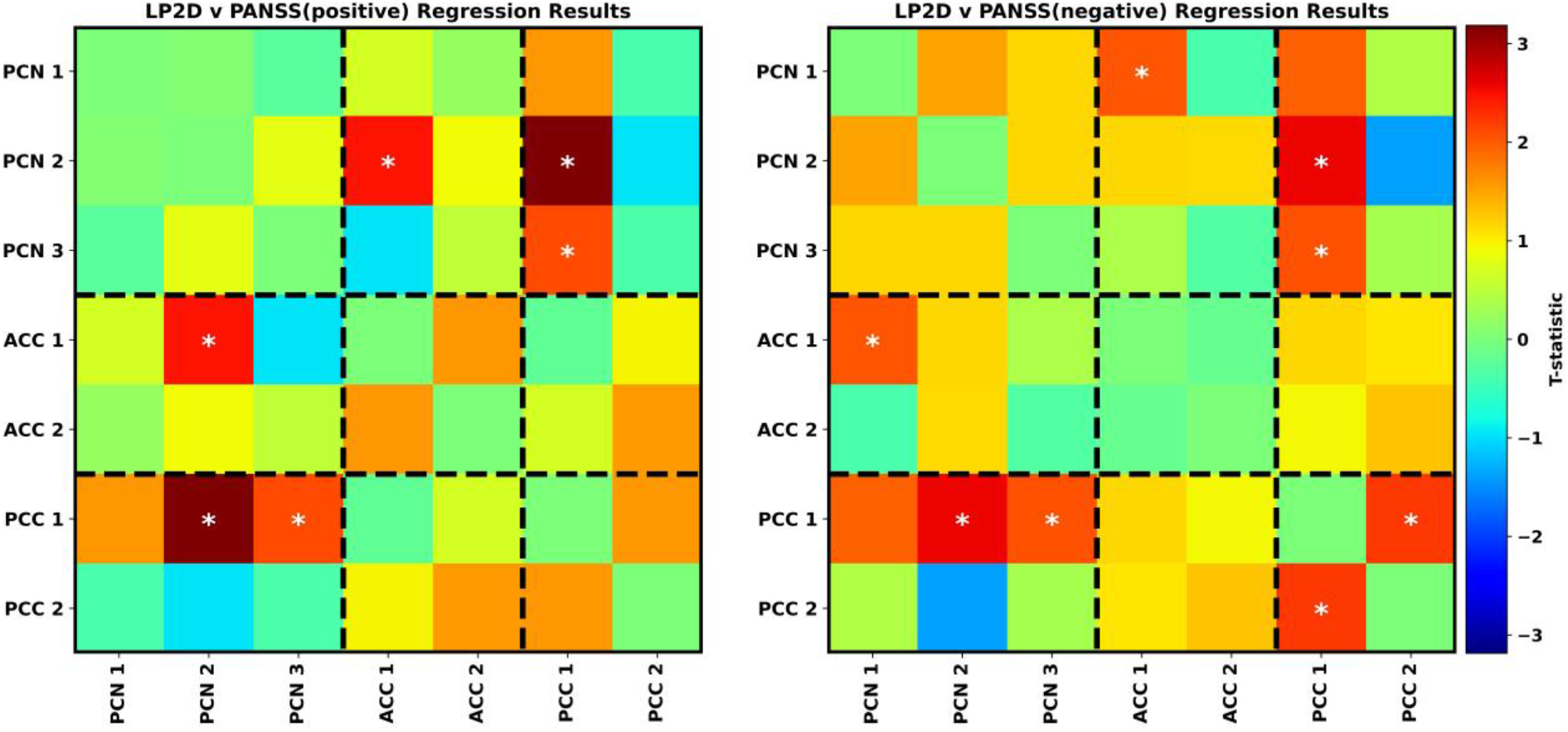
Relationship between Symptom Severity and LP2D Stability Features. The lefthand and righthand figures show LP2D relationships with positive and negative PANSS scores, respectively. Heatmaps indicate t-statistic values resulting from the regression analysis, and both panels share the same color bar to the right of the figure. Results are arranged in the form of a connectivity matrix for ease of interpretation, with the corresponding brain regions shown on the x- and y-axes. Domain pairs are separated by black dashed lines, and white asterisks indicate features with a significant uncorrected correlation with symptom severity.

## DISCUSSION

The goals of this study were (1) to present a framework for dFNC state analysis that accounts for the inherent variability of data within states and (2) provide additional insights into the effects of SZ upon DMN dynamics. We identified 5 fuzzy states of DMN activity, characterized the states using a novel explainability approach, identified effects of SZ upon DMN dynamics and state stability, and identified relationships between state stability and symptom severity.

Interestingly, the centroids for the 5 fuzzy states that we identified diverge greatly from those identified in previous studies [13]. States 0, 1, 3, and 4 domain pairs (e.g., PCN/PCN, PCN/ACC) were much more uniform within states relative to those identified in [13]. For example, in state 1 ACC/PCN and ACC/PCC were highly negative while all other domain pairs were slightly positive. Only state 2, in which HCs spent more time, and state 0, to a much smaller degree, were similar to the state centroids identified in [13] in their varied dFNC across domain pairs.

HCs spent much more time in state 2, a state characterized by having varying magnitudes of both positive and negative dFNC. Within state 2, HCs significantly higher levels of variance and entropy than SZs, and relative to SZ state 0 entropy and variance, HCs had higher levels of state 2 variance and entropy. In contrast, SZs spent much more time in state 0, a state with highly negative intra-ACC activity and varied activity in other domain pairs. Intra-ACC activity was highly important to SZ dynamics, as they lacked stability to perturbation of intra-ACC activity. There have historically been contradictory findings related to the effects of SZ upon the ACC. However, our findings fit with those of multiple studies [13][27]. State 0 also had higher intra-PCN activity than state 2, which conforms to previous studies that have found higher intra-PCN activity in SZs than HCs [31]. SZs also tended to be less stable to PCC2/PCN perturbation, whereas HCs tended to be less stable than SZs to PCC1/PCN perturbation. Although some studies agree with our finding that SZs spend more time in a state with more positive connectivity [31] (i.e., state 0 versus state 2), many previous studies have found SZ inter-domain connectivity to be less than HCs [10]. This difference in results is likely attributable to the use of different regions within the DMN [10][64], the use of whole-brain analyses [64] that have been shown to obscure the activity of the DMN [13], [23], or the use of multiple disorders that can disguise the effects of SZ [12].

HCs tended to spend slightly more time or a comparable amount of time in states outside of state 2 than SZs spent out of state 0. Additionally, based on our state correlation features, they tended to shift their similarity to more fuzzy state centroids than SZs when transitioning from state 2 (i.e., to all 4 states rather than only 3 other states). Additionally, while SZs tended to have higher KLD feature values than HCs (i.e., mean, max, range, SD, and start/stop), most of our KLD dynamical features are likely to capture temporally localized information than more distributed information. While SZs had slightly higher mean KLD over time than HCs based on visual assessment of distributions, there was no noticeable difference statistically or otherwise between SZ and HC median KLD, which supports the presence of extreme KLD SZ values that would greatly influence our mean, max, range, and SD KLD features. Our KLD findings could indicate that the aberrant SZ dynamics vary in a more temporally localized manner than healthy dynamics [24] and paired with our correlation findings indicate that dynamics over time could potentially be reduced in SZ relative to HCs [29].

Several studies have previously identified relationships between DMN dFNC features and symptom severity [12], [13]. Relative to features extracted in previous studies [13], we found that our use of LP2D to assess the stability of samples to perturbation of dFNC features was uncovered a large number of relationships with both positive and negative symptom severity. Accounting for age and gender, PCC1/PCN and ACC1/PCN stability was highly related to both positive and negative symptom severity. Previous studies have related PCC/PCN connectivity with positive symptom severity [31] and the PCN to negative symptom severity [34]. Additionally, intra-PCC stability was also related to negative symptom severity [10].

While our findings hold great significance for the domain of SZ analysis, our proposed explainable fuzzy clustering framework has broader implications. Our use of fuzzy clustering enables insight not only into inter-state dynamics but also into intra-state dynamics. Furthermore, our approach accounts for the inherent variation in how similar samples are to their assigned centroid and to samples assigned to other centroids. These challenges will be present in any dFNC clustering analysis. Our dynamical and stability features also represent key advances, providing new insights into disorder-related activity. Additionally, as we showed by thresholding the state probabilities and calculating OCR and NST values, traditional k-means-based features can still be extracted with fuzzy c-means. That paired with the ease of implementation of fuzzy c-means using existing Python packages [52] and MATLAB [53] means that our approach could easily replace existing dFNC k-means clustering approaches in future studies.

The explainability methods that we present also represent advances for the field, providing a quantitative estimate of the importance of each dFNC feature to the clustering. Relative to G2PC, our approach accounts for the effects of perturbation upon 100% of samples. Additionally, different metrics can be applied to the resulting KLD values to gain different insights into the effects of perturbation or to produce more or less sparse explanations (e.g., in total versus median KLD).

### Limitations and Future Opportunities

Our study and approach have several limitations. Namely, we used a 40-second window when calculating dFNC, and while studies have shown that to be a reasonable window size [56], the window size can affect the dynamics and findings. Fuzzy c-means has a couple limitations. (1) Similar to k-means clustering, it is affected by outliers. However, there are variations of fuzzy c-means that are capable of addressing this problem [65]. Additionally, fuzzy c-means can be more computationally intensive than k-means clustering [66]. However, an approach similar to iSparse k-means could be easily adapted to work with fuzzy c-means [67]. Lastly, future research directions might include analyzing the reproducibility of our results across datasets similar to [13] or analyzing multiple disorders in a single analysis like [4].

## CONCLUSION

The analysis of rs-fMRI dFNC data using hard clustering methods to identify states that summarize brain dynamics is a common analysis approach that has provided insights into many neurological and neuropsychiatric disorders. However, the use of hard clustering approaches (e.g., k-means clustering) can obscure key information related to how similar samples are to their respective centroids or to samples assigned to other cluster centroids. As such, the use of hard clustering could obscure disorder-relevant dynamics. In this study, we present a novel explainable fuzzy clustering framework. We present 7 new types of dynamical features and sample stability-based features that provide unique insights into brain dynamics while also demonstrating how traditional dynamical features used in hard clustering analyses can also be included in our analysis. Lastly, we present two novel explainability approaches that help characterize the fuzzy states identified using our clustering approach. We demonstrate our framework within the context of SZ DMN analysis, identifying aberrant dynamics in SZs and uncovering relationships between SZ symptom severity and precuneus interactions with the anterior cingulate cortex and posterior cingulate cortex. Our framework provides greater insight into disease dynamics than traditional hard clustering approaches. Furthermore, it can be implemented with an ease comparable to the standard k-means clustering approach using existing code packages. As such, it represents an ideal method for future widespread use in rs-fMRI dFNC analysis and could lead to an improved understanding of the effects of many neurological and neuropsychological disorders upon brain dynamics.

## ACKNOWLEDGMENTS

We thank those who collected the FBIRN dataset. This research is supported by NIH R01MH123610, NIH R01MH118695, and NSF 2112455.

## REFERENCES

[1] M. S. E. Sendi et al., “The link between brain functional network connectivity and genetic risk of Alzheimer’s disease,” bioRxiv, 2021, doi: 10.1002/alz.050101.

[2] E. Damaraju et al., “Dynamic functional connectivity analysis reveals transient states of dysconnectivity in schizophrenia,” NeuroImage Clin., vol. 5, no. July, pp. 298–308, 2014, doi: 10.1016/j.nicl.2014.07.003.

[3] E. Zendehrouh et al., “Aberrant Functional Network Connectivity Transition Probability in Major Depressive Disorder,” in 42nd Annual International Conference of the IEEE Engineering in Medicine and Biology Society (EMBC), 2020, pp. 1493–1496.

[4] X. jie Wu et al., “Functional network connectivity alterations in schizophrenia and depression,” Psychiatry Res. - Neuroimaging, vol. 263, no. August 2016, pp. 113–120, 2017, doi: 10.1016/j.pscychresns.2017.03.012.

[5] H. Dini et al., “Dynamic Functional Connectivity Predicts Treatment Response to Electroconvulsive Therapy in Major Depressive Disorder,” Front. Hum. Neurosci., vol. 15, no. July, pp. 1–11, 2021, doi: 10.3389/fnhum.2021.689488.

[6] J. Y. Chun, M. S. E. Sendi, J. Sui, D. Zhi, and V. D. Calhoun, “Visualizing Functional Network Connectivity Difference between Healthy Control and Major Depressive Disorder Using an Explainable Machine-learning Method,” in 2020 42nd Annual International Conference of the IEEE Engineering in Medicine & Biology Society (EMBC), 2020, pp. 955–960, doi: 10.1109/BIBE50027.2020.00162.

[7] C. A. Ellis, M. L. Sancho, R. Miller, and V. Calhoun, “Exploring Relationships between Functional Network Connectivity and Cognition with an Explainable Clustering Approach,” in 2022 IEEE 22nd International Conference on Bioinformatics and Bioengineering (BIBE), 2022, pp. 23–26, doi: 10.1109/BIBE55377.2022.00066.

[8] M. S. E. Sendi, C. A. Ellis, R. L. Milller, D. H. Salat, and V. D. Calhoun, “The relationship between dynamic functional network connectivity and spatial orientation in healthy young adults,” bioRxiv, 2021.

[9] L. Sanfratello, J. Houck, and V. D. Calhoun, “Dynamic Functional Network Connectivity In Schizophrenia With MEG And fMRI, Do Different Time Scales Tell A Different Story?,” Brain Connect., 2019, doi: 10.1089/brain.2018.0608.

[10] Y. Du et al., “Interaction among subsystems within default mode network diminished in schizophrenia patients: A dynamic connectivity approach,” Schizophr. Res., vol. 170, no. 1, pp. 55–65, 2016, doi: 10.1016/j.schres.2015.11.021.

[11] R. L. Miller et al., “Higher dimensional meta-state analysis reveals reduced resting fMRI connectivity dynamism in schizophrenia patients,” PLoS One, vol. 11, no. 3, pp. 1–24, 2016, doi: 10.1371/journal.pone.0149849.

[12] M. S. E. Sendi, H. Dini, L. E. Bruni, and V. D. Calhoun, “Default mode network dynamic functional network connectivity predicts psychotic symptom severity,” Proc. Annu. Int. Conf. IEEE Eng. Med. Biol. Soc. EMBS, vol. 2022-July, pp. 247–250, 2022, doi: 10.1109/EMBC48229.2022.9871542.

[13] M. S. E. Sendi et al., “Aberrant Dynamic Functional Connectivity of Default Mode Network in Schizophrenia and Links to Symptom Severity,” Front. Neural Circuits, vol. 15, no. March, pp. 1–14, 2021, doi: 10.3389/fncir.2021.649417.

[14] M. S. Salman, Y. Du, and V. D. Calhoun, “Identifying FMRI dynamic connectivity states using affinity propagation clustering method: Application to schizophrenia,” ICASSP, IEEE Int. Conf. Acoust. Speech Signal Process. - Proc., pp. 904–908, 2017, doi: 10.1109/ICASSP.2017.7952287.

[15] A. Abrol et al., “Replicability of time-varying connectivity patterns in large resting state fMRI samples,” Neuroimage, vol. 163, pp. 160–176, 2017, doi: 10.1016/j.neuroimage.2017.09.020.Replicability.

[16] C. A. Ellis, R. L. Miller, and V. D. Calhoun, “A Convolutional Autoencoder-based Explainable Clustering Approach for Resting-State EEG Analysis,” in bioRxiv, 2023, pp. 3–6.

[17] C. A. Ellis, A. Sattiraju, R. Miller, and V. Calhoun, “Examining Effects of Schizophrenia on EEG with Explainable Deep Learning Models,” 2022.

[18] C. A. Ellis, A. Sattiraju, R. Miller, and V. Calhoun, “Examining Reproducibility of EEG Schizophrenia Biomarkers Across Explainable Machine Learning Models,” in 2022 IEEE 22nd International Conference on Bioinformatics and Bioengineering (BIBE), 2022, pp. 305–308, doi: 10.1109/BIBE55377.2022.00069.

[19] T. J. Gawne et al., “A multimodal magnetoencephalography 7 T fMRI and 7 T proton MR spectroscopy study in first episode psychosis,” npj Schizophr., vol. 6, no. 1, pp. 1–9, 2020, doi: 10.1038/s41537-020-00113-4.

[20] W. Yan et al., “Discriminating schizophrenia using recurrent neural network applied on time courses of multi-site FMRI data,” EBioMedicine, vol. 47, pp. 543–552, Sep. 2019, doi: 10.1016/j.ebiom.2019.08.023.

[21] C. A. Ellis, R. L. Miller, and V. D. Calhoun, “An Approach for Estimating Explanation Uncertainty in fMRI dFNC Classification,” 2022 IEEE 22nd Int. Conf. Bioinforma. Bioeng., 2022.

[22] C. A. Ellis, R. L. Miller, and V. D. Calhoun, “Towards Greater Neuroimaging Classification Transparency via the Integration of Explainability Methods and Confidence Estimation Approaches,” Informatics Med. Unlocked, vol. 37, 2023, doi: https://doi.org/10.1016/j.imu.2023.101176.

[23] C. A. Ellis, M. S. E. Sendi, R. L. Miller, and V. D. Calhoun, “An Unsupervised Feature Learning Approach for Elucidating Hidden Dynamics in rs-fMRI Functional Network Connectivity,” in 2022 44th Annual International Conference of the IEEE Engineering in Medicine & Biology Society (EMBC), 2022, pp. 4449–4452.

[24] M. Rahman et al., “Interpreting models interpreting brain dynamics,” Sci. Rep., vol. 12, pp. 1–16, 2022.

[25] V. Bliksted et al., “Hyper- and Hypomentalizing in Patients with First-Episode Schizophrenia: FMRI and Behavioral Studies,” Schizophr. Bull., vol. 45, no. 2, pp. 377–385, 2019, doi: 10.1093/schbul/sby027.

[26] S. J. H. Ebisch et al., “Disrupted relationship between ‘resting state’ connectivity and task-evoked activity during social perception in schizophrenia,” Schizophr. Res., vol. 193, pp. 370–376, 2018, doi: 10.1016/j.schres.2017.07.020.

[27] D. K. Shukla et al., “Anterior cingulate glutamate and GABA associations on functional connectivity in schizophrenia,” Schizophr. Bull., vol. 45, no. 3, pp. 647–658, 2019, doi: 10.1093/schbul/sby075.

[28] S. M. Hare et al., “Salience-default mode functional network connectivity linked to positive and negative symptoms of schizophrenia,” Schizophr. Bull., vol. 45, no. 4, pp. 892–901, 2019, doi: 10.1093/schbul/sby112.

[29] A. Kottaram et al., “Brain network dynamics in schizophrenia: Reduced dynamism of the default mode network,” Hum. Brain Mapp., vol. 40, no. 7, pp. 2212–2228, 2019, doi: 10.1002/hbm.24519.

[30] G. B. Chand, D. S. Thakuri, B. Soni, and S. Kingshighway Blvd St Louis, “Disrupted controlling mechanism of salience network on default-mode network and central-executive network in schizophrenia,” bioRxiv, pp. 1–19, 2021, [Online]. Available: https://doi.org/10.1101/2021.12.03.471183.

[31] S. Whitfield-Gabrieli et al., “Hyperactivity and hyperconnectivity of the default network in schizophrenia and in first-degree relatives of persons with schizophrenia,” Proc. Natl. Acad. Sci. U. S. A., vol. 106, no. 4, pp. 1279–1284, 2009, doi: 10.1073/pnas.0809141106.

[32] Q. Yu and V. D. Calhoun, “Resting-State Functional Network Disturbances in Schizophrenia,” Brain Netw. Dysfunct. Neuropsychiatr. Illn., pp. 187–215, 2021, doi: 10.1007/978-3-030-59797-9_10.

[33] B. Sen, B. Mueller, B. Klimes-Dougan, K. Cullen, and K. K. Parhi, “Classification of Major Depressive Disorder from Resting-State fMRI,” Proc. Annu. Int. Conf. IEEE Eng. Med. Biol. Soc. EMBS, no. Mdd, pp. 3511–3514, 2019, doi: 10.1109/EMBC.2019.8856453.

[34] Y. Sun, S. L. Collinson, J. Suckling, and K. Sim, “Dynamic reorganization of functional connectivity reveals abnormal temporal efficiency in schizophrenia,” Schizophr. Bull., vol. 45, no. 3, pp. 659–669, 2019, doi: 10.1093/schbul/sby077.

[35] M. S. E. Sendi et al., “Disrupted Dynamic Functional Network Connectivity Among Cognitive Control Networks in the Progression of Alzheimer’s Disease,” Brain Connect., pp. 1–25, 2021, doi: 10.1089/brain.2020.0847.

[36] M. S. E. Sendi et al., “Alzheimer’s Disease Projection From Normal to Mild Dementia Reflected in Functional Network Connectivity: A Longitudinal Study,” Front. Neural Circuits, vol. 14, no. January, 2021, doi: 10.3389/fncir.2020.593263.

[37] M. S. E. Sendi et al., “Multiple overlapping dynamic patterns of the visual sensory network in schizophrenia,” Schizophr. Res., vol. 228, pp. 103–111, 2021, doi: 10.1016/j.schres.2020.11.055.

[38] Y. Du et al., “NeuroMark: An automated and adaptive ICA based pipeline to identify reproducible fMRI markers of brain disorders,” NeuroImage Clin., vol. 28, no. August, p. 102375, 2020, doi: 10.1016/j.nicl.2020.102375.

[39] C. A. Ellis, R. L. Miller, and V. D. Calhoun, “Neuropsychiatric Disorder Subtyping Via Clustered Deep Learning Classifier Explanations,” in bioRxiv, 2022, pp. 12–15.

[40] Z. Fu et al., “Dynamic functional network reconfiguration underlying the pathophysiology of schizophrenia and autism spectrum disorder,” Hum. Brain Mapp., vol. 42, no. 1, pp. 80–94, 2021, doi: 10.1002/hbm.25205.

[41] M. S. E. Sendi, J. Y. Chun, and V. D. Calhoun, “Visualizing functional network connectivity difference between middle adult and older subjects using an explainable machine-learning method,” in Proceedings - IEEE 20th International Conference on Bioinformatics and Bioengineering, BIBE 2020, 2020, pp. 955–960, doi: 10.1109/BIBE50027.2020.00162.

[42] Z. Fu et al., “Dynamic state with covarying brain activity-connectivity: On the pathophysiology of schizophrenia,” Neuroimage, vol. 224, no. July 2020, p. 117385, 2021, doi: 10.1016/j.neuroimage.2020.117385.

[43] C. A. . Ellis, R. L. . Miller, and V. D. . Calhoun, “Identifying Neuropsychiatric Disorder Subtypes and Subtype-Dependent Variation in Diagnostic Deep Learning Classifier Performance,” bioRxiv, pp. 2–5, 2022.

[44] M. J, “Some Methods for Classification and Analysis of MultiVariate Observations,” Proc Berkeley Symp. Math. Stat. Probab., vol. 5, no. 1, pp. 281–297, 1965.

[45] F. Pedregosa, R. Weiss, and M. Brucher, “Scikit-learn: Machine Learning in Python,” J. Mach. Learn. Res., vol. 12, pp. 2825–2830, 2011.

[46] E. A. Allen, E. Damaraju, S. M. Plis, E. B. Erhardt, T. Eichele, and V. D. Calhoun, “Tracking whole-brain connectivity dynamics in the resting state,” Cereb. Cortex, vol. 24, no. 3, pp. 663–676, 2014, doi: 10.1093/cercor/bhs352.

[47] C. A. Ellis, M. S. E. Sendi, E. P. T. Geenjaar, S. M. Plis, R. L. Miller, and V. D. Calhoun, “Algorithm-Agnostic Explainability for Unsupervised Clustering,” pp. 1–22, 2021, [Online]. Available: http://arxiv.org/abs/2105.08053.

[48] R. Suganya and R. Shanthi, “Fuzzy C-Means Algorithm-A Review,” Int. J. Sci. Res. Publ., vol. 2, no. 11, pp. 2250–3153, 2012, [Online]. Available: https://www.ijsrp.org.

[49] S. Demircan and H. Kahramanli, “Application of fuzzy C-means clustering algorithm to spectral features for emotion classification from speech,” Neural Comput. Appl., vol. 29, no. 8, pp. 59–66, 2018, doi: 10.1007/s00521-016-2712-y.

[50] S. Munusamy and P. Murugesan, “Modified dynamic fuzzy c-means clustering algorithm – Application in dynamic customer segmentation,” Appl. Intell., vol. 50, no. 6, pp. 1922–1942, 2020, doi: 10.1007/s10489-019-01626-x.

[51] S. Kahali, J. K. Sing, and P. K. Saha, “A new entropy-based approach for fuzzy c-means clustering and its application to brain MR image segmentation,” Soft Comput., vol. 23, no. 20, pp. 10407–10414, 2019, doi: 10.1007/s00500-018-3594-y.

[52] “Scikit-Fuzzy,” 2022. https://scikit-fuzzy.github.io/scikit-fuzzy/.

[53] “Fuzzy C-Means Clustering,” MATLAB R2022b. https://www.mathworks.com/help/fuzzy/fuzzy-c-means-clustering.html.

[54] T. G. M. van Erp et al., “Neuropsychological profile in adult schizophrenia measured with the CMINDS,” Psychiatry Res., vol. 230, no. 3, pp. 826–834, 2015, doi: 10.1016/j.psychres.2015.10.028.Neuropsychological.

[55] S. R. Kay, A. Fiszbein, and L. A. Opler, “The positive and negative syndrome scale (PANSS) for schizophrenia,” Schizophr. Bull., vol. 13, no. 2, pp. 261–276, 1987, doi: 10.1093/schbul/13.2.261.

[56] Z. Fu, Y. Du, and V. D. Calhoun, “The Dynamic Functional Network Connectivity Analysis Framework,” Engineering, vol. 5, no. 2, pp. 190–193, 2019, doi: 10.1016/j.eng.2018.10.001.

[57] Y. Du et al., “Dynamic functional connectivity impairments in early schizophrenia and clinical high-risk for psychosis,” Neuroimage, vol. 180, pp. 632–645, 2018, doi: 10.1016/j.neuroimage.2017.10.022.Dynamic.

[58] L. E. O. Breiman, “Random Forests,” Mach. Learn., vol. 45, pp. 5–32, 2001.

[59] A. Fisher, C. Rudin, and F. Dominici, “Model Class Reliance: Variable Importance Measures for any Machine Learning Model Class, from the ‘Rashomon’ Perspective,” arXiv Prepr. arXiv 1801.01489v1, 018.

[60] Y. Kakizawa, “Discriminant analysis for non-gaussian vector stationary processes,” J. Nonparametr. Stat., vol. 7, no. 2, pp. 187–203, 1996, doi: https://doi.org/10.1080/10485259608832698.

[61] M. G. Kendall, “A New Measure of Rank Correlation,” Biometrika, vol. 30, no. 1/2, pp. 81–93, 1938.

[62] P. Virtanen et al., “SciPy 1.0: fundamental algorithms for scientific computing in Python,” Nat. Methods, vol. 17, no. 3, pp. 261–272, 2020, doi: 10.1038/s41592-019-0686-2.

[63] Y. Benjamini and Y. Hochberg, “Controlling the False Discovery Rate: A Practical and Powerful Approach to Multiple Testing,” J. R. Stat. Soc. Ser. B, vol. 57, no. 1, pp. 289–300, 1995, doi: 10.1111/j.2517-6161.1995.tb02031.x.

[64] S. Li et al., “Dysconnectivity of multiple brain networks in schizophrenia: A meta-analysis of resting-state functional connectivity,” Front. Psychiatry, vol. 10, no. JULY, pp. 1–11, 2019, doi: 10.3389/fpsyt.2019.00482.

[65] S. Askari, “Fuzzy C-Means clustering algorithm for data with unequal cluster sizes and contaminated with noise and outliers: Review and development,” Expert Syst. Appl., vol. 165, no. March 2020, p. 113856, 2021, doi: 10.1016/j.eswa.2020.113856.

[66] S. Ghosh and S. K. Dubey, “A Comparative Analysis of Fuzzy C-Means Clustering and K Means Clustering Algorithms,” Int. J. Adv. Comput. Sci. Appl., vol. 4, no. 4, 2013.

[67] M. S. E. Sendi, D. H. Salat, R. L. Miller, and V. D. Calhoun, “Two-step clustering-based pipeline for big dynamic functional network connectivity data,” Front. Neurosci., vol. 16, 2022, doi: 10.3389/fnins.2022.895637.

