## Supplementary Figures for "Explainable Fuzzy Clustering Framework Reveals Divergent Default Mode Network Connectivity Dynamics in Schizophrenia"

**Supplementary Materials**

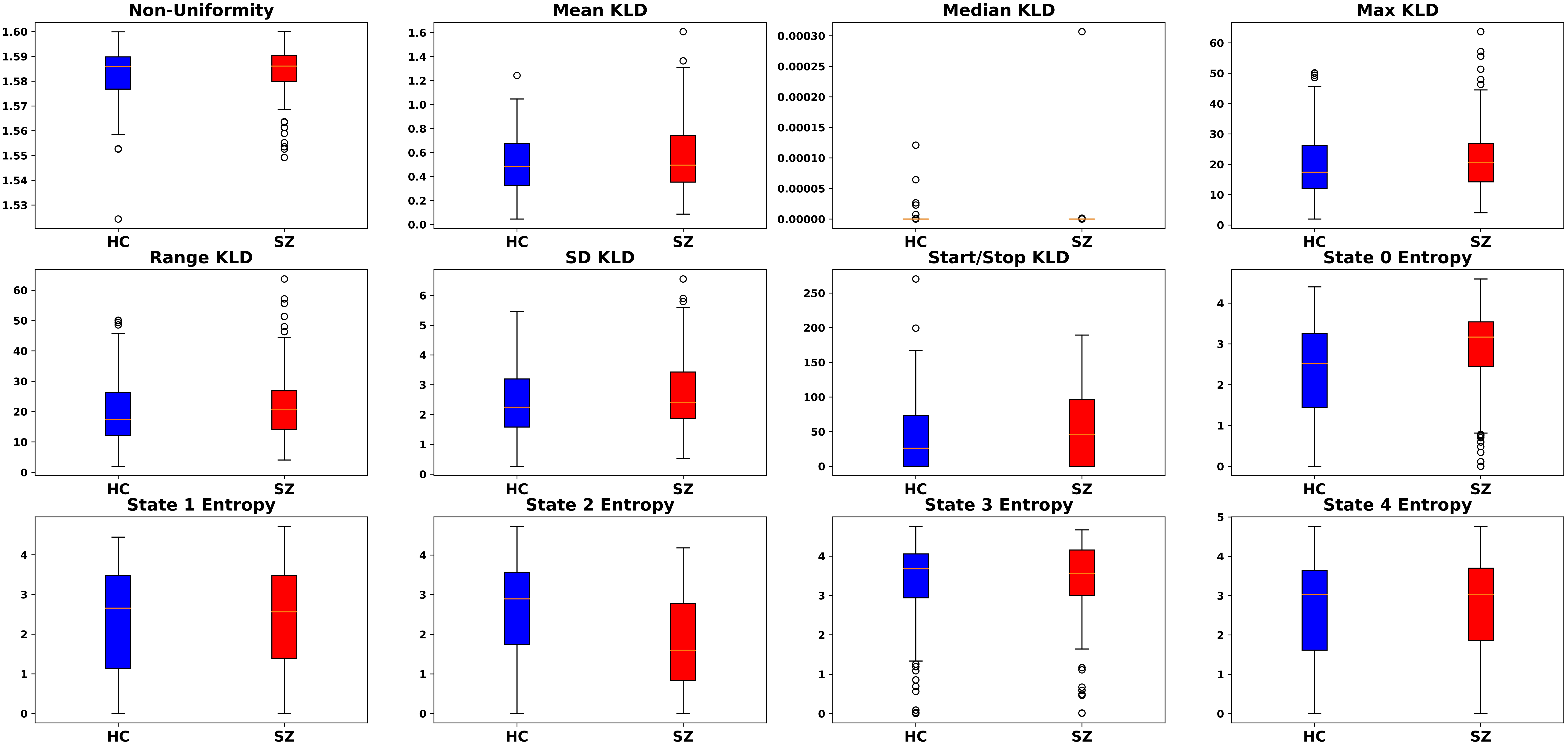

Supplementary Figure 1. Boxplots of Non-Uniformity, KLD, and Entropy Features. Each panel shows boxplots for the dynamical features for HCs (left) and SZs (right). The title of each panel indicates the dynamical feature included in the boxplots.

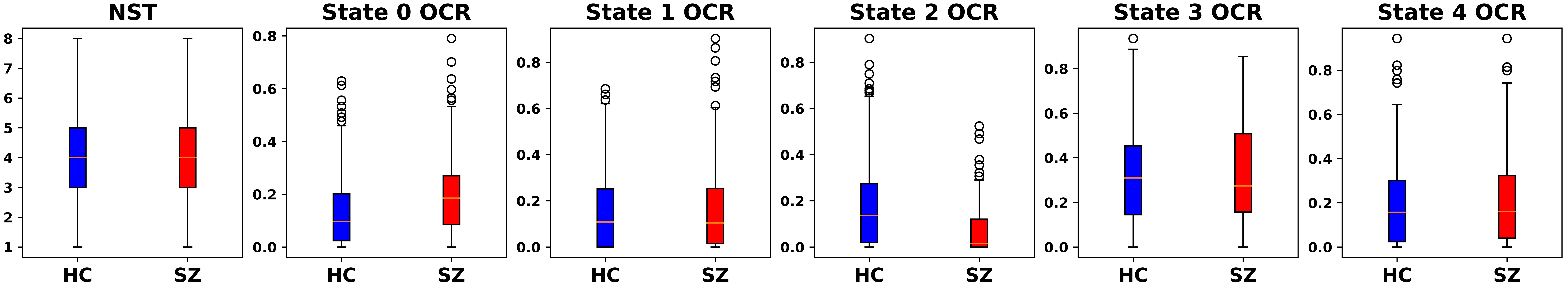

Supplementary Figure 2. Traditional NST + OCR Features. Each panel shows boxplots for the dynamical features for HCs (left) and SZs (right). The title of each panel indicates the dynamical feature included in the boxplots.

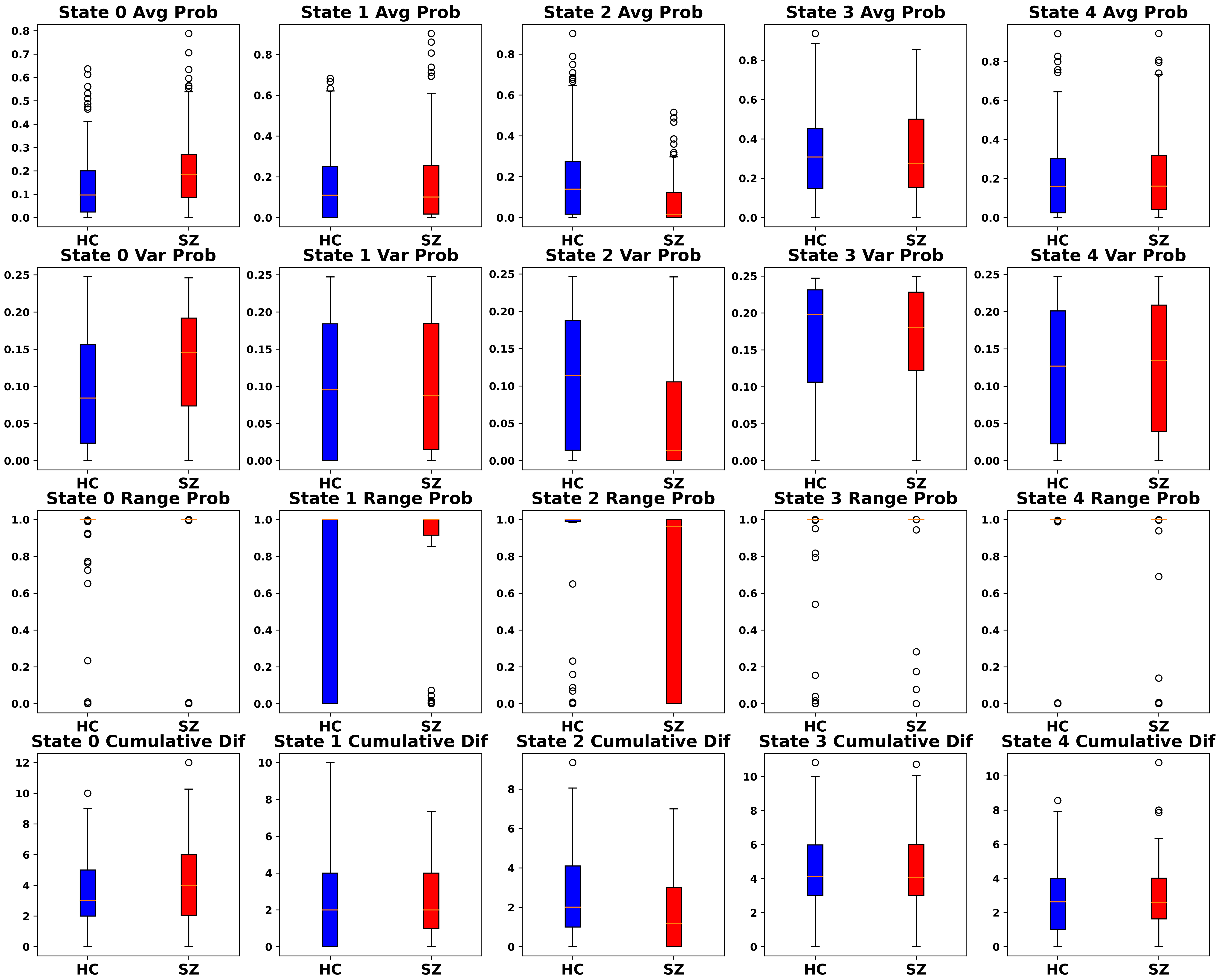

Supplementary Figure 3. Boxplots of Average, Variance, Range, and Cumulative Difference Features. Each panel shows boxplots for the dynamical features for HCs (left) and SZs (right). The title of each panel indicates the dynamical feature included in the boxplots.

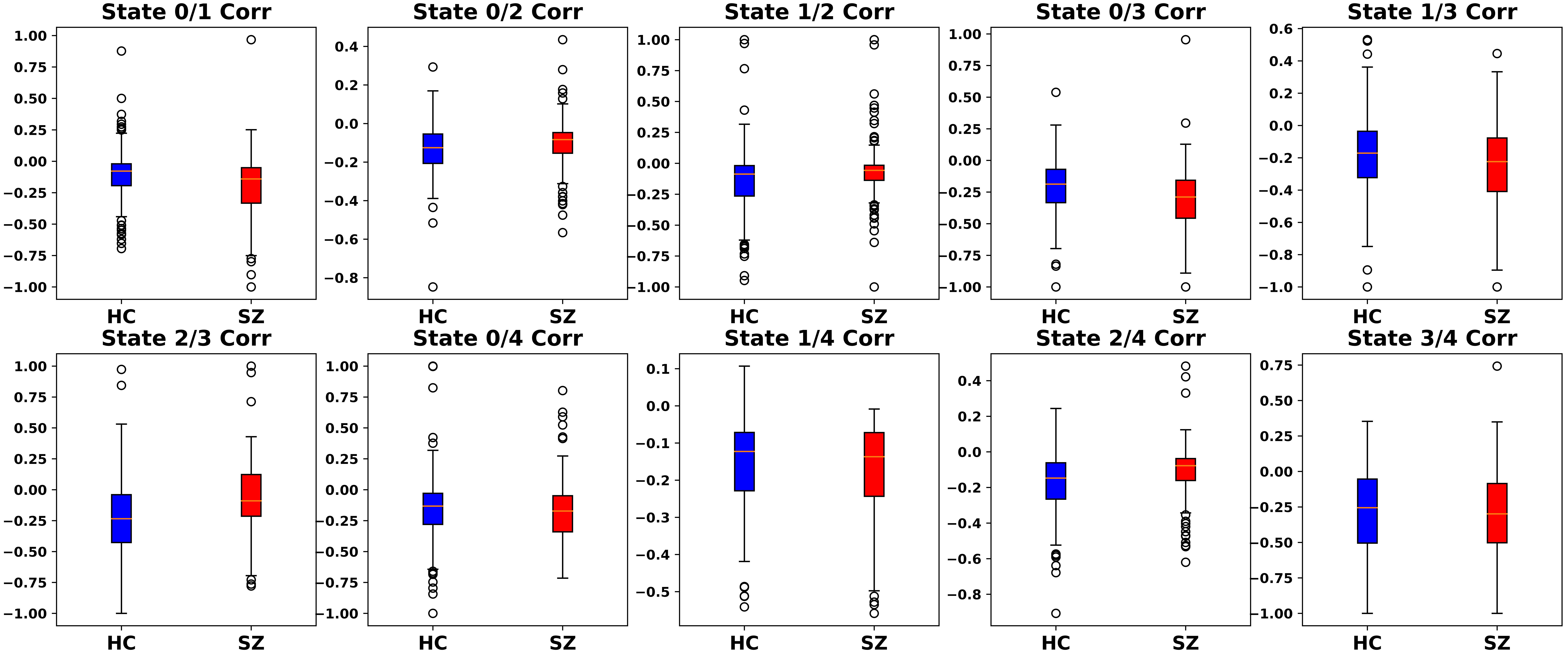

Supplementary Figure 4. Boxplots of Correlation Features. Each panel shows boxplots for the dynamical features for HCs (left) and SZs (right). The title of each panel indicates the dynamical feature included in the boxplots.
